# A VERSATILE LIBRARY OF TETRACYCLINE-INDUCIBLE AND REPRESSIBLE VECTORS FOR FINE-TUNED GENE EXPRESSION

**DOI:** 10.64898/2026.04.30.721401

**Authors:** Akshay V Nair, Shinto James, Vikas Jain

## Abstract

The genus *Mycobacterium* is increasingly recognized as a major clinical concern due to diseases such as tuberculosis, along with the emergence of antimicrobial-resistant strains, underscoring the urgent need for advanced genetic tools to study mycobacterial biology and pathogenesis. Progress in this area relies heavily on the functional characterization of previously unannotated genes, which necessitates tightly regulated expression systems. Here, we report the development of an improved tetracycline-regulated vector platform, comprising the episomal pM(R)T2 and integrative pMI(R)T2 series, which builds upon the previously described pMT vector system. The ‘T2’ vector series incorporates a fine-tuned TetRO system for enhanced transcriptional control. The pMT2 vectors function as tetracycline-inducible systems, whereas the pMRT2 variants utilize a reverse tetracycline repressor (RevTetR) to enable tetracycline-repressible gene regulation. Additionally, the integrative variant, pMI(R)T2 switches the *oriM* element with the integrase and *attP* sites derived from mycobacteriophage L5, facilitating stable genomic integration and controlled expression of concentration-sensitive genes, including toxins. To expand the selection flexibility, the pAN(R)Tet series replaces the kanamycin resistance cassette with a hygromycin resistance cassette. Functional validation of gene regulation in *M. smegmatis* and *M. bovis* BCG shows that both TetR and RevTetR systems provide reliable inducible and repressible controls, respectively, upon anhydrotetracycline addition. Taken together, these vectors constitute a versatile, tightly regulated genetic toolkit with significant potential to accelerate research and therapeutic development in mycobacterial systems.

## INTRODUCTION

The Mycobacterium genus is a diverse group of species that includes many major human pathogens, such as Mycobacterium tuberculosis and Mycobacterium leprae, which collectively impose a significant global health burden. Tuberculosis caused by the *Mycobacterium tuberculosis (Mtb)* has killed around 1.3 million people globally, with an estimated 1.7 billion people latently infected. Co-morbidities like diabetes, HIV, and COVID-19, along with the emergence of drug-resistant strains, have further increased the mortality of tuberculosis infections, making it a global threat**(WHO 2023, n.d.)**. What makes treatment difficult is its remarkable ability to adapt to its environment upon infection, evade the host immune response, and survive harsh lung microenvironments. The resilience of *Mtb* can be attributed to its thick, unusually lipid-rich cell wall, which creates a physical barrier against various antibiotics and toxic molecules **(Alderwick et al., 2015)**. Other factors, such as bacterial persistence and high mutation rates, also contribute to the acquisition of antibiotic resistance genes, while the ability to form biofilms further enhances its resilience **(Judd et al., 2021; Sparks et al., 2023; Yang et al., 2024)**.

Plasmids are mobile genetic elements that play an important role in bacterial biology, from helping in horizontal gene transfer and conferring antibiotic resistance to driving the evolution of virulence mechanisms **(Krawczyk et al., 2018)**. Despite their importance, the diversity and functional complexity of plasmids, especially in mycobacteria remain underexplored, partly due to historical limitations in sequencing technologies and poor efficiency of plasmid transformation **(Snapper et al., 1990)**. Although, Recent advancements in genetic tools and long-read sequencing have begun to shed light onto the dynamic nature of plasmid architecture and their contributions to mycobacterial adaptation, yet a systematic resource for studying these elements remains to be desired **(Dumas et al., 2016; Xu et al., 2024)**.

The development of efficient plasmid transformation (ept) systems in *M. smegmatis*, particularly the eptC mutant strain mc^2^155, has revolutionized mycobacterial genetics by providing efficient uptake and stable plasmid maintenance. This breakthrough has facilitated the construction of early shuttle phasmids and hybrid plasmids, such as pAL5000-based vectors, which can replicate in both *E. coli* and mycobacteria **(Panas et al., 2014; Snapper et al., 1990; Sparks et al., 2023)**. Subsequent innovations, including temperature-sensitive origins of replication **(Guilhot et al., 1992; Pelicic et al., 1997)** and site-specific integration systems **(Saviola & Bishai, 2004)**, have further expanded the toolkit for plasmid engineering.

Of the various expression systems developed for mycobacteria, the tetracycline-regulation system is widely used owing to its tight regulation, fast and dose-dependent activity **(Blokpoel et al., 2005; Carroll et al., 2005; Seniya et al., 2020)**, presence of both inducible (TetON) and repressible (TetOFF) systems, use of multiple effectors and compatibility with different mycobacterial species and growth conditions **(Schnappinger & Ehrt, 2014)**.

The tetracycline-inducible system, also called the TetON system, comprises two divergently arranged promoters, one regulates the expression of the tetracycline repressor (TetR), while the other controls the target gene. These promoters include two operator (TetO) sites bound by TetR in the absence of tetracycline, inhibiting gene expression. Upon the addition of tetracycline, TetR binds to tetracycline and undergoes a conformational change that causes it to disassociate from TetO and allow downstream gene expression **(Carroll et al., 2005)**. Furthermore, the functional and structural characterization of the Tet repressor led to the development of the tetracycline-repressible system or the TetOFF system. They use tetracycline and its derivatives as a co-repressor to shut off the expression of downstream genes in their presence. TetOFF is comprised of multiple systems based on their mode of gene repression.

TRE (tet responsive elements), PipOFF, RevTetR **(Bertram et al., 2022 PMID 34713957)**.

The pMT vector library, previously reported by our lab, contains the tetracycline promoter-repressor cassette derived from *Corynebacterium glutamicum*. Despite modifications to minimize basal expression, the promoter exhibits considerable leaky expression in *Mycobacterium smegmatis* **(Seniya et al., 2020)**. Among the three tetracycline-inducible systems introduced into mycobacterial species, the *E. coli* Tet system has shown the lowest level of leaky expression and the highest fold induction upon inducer exposure **(Meijers et al., 2020; Schnappinger & Ehrt, 2014)**.

As robust regulation is paramount for the study of different genes in mycobacteria, we, in this study, have designed a new library of vectors pMT2, which is based on the previously described pMT vector having the more regulated E. coli Tet system. We have further developed a repressible vector library, pMRT2, derived from pMT2 as part of the TetON and OFF system. pMRT2 uses a modified tet repressor that shows a reverse phenotype, called the reverse Tet repressor **(Klotzsche et al., 2009)**. The tet repressor contains a single amino acid mutation (V99E) that converts it into the reverse tet repressor (RevTetR). The valine residue at the 99th position of the repressor protein lies within its hydrophobic core region. Introducing a charged residue, such as glutamic acid, destabilizes the core, shifting the H-bonding network between helices 1 and 6, leading to a conformational rearrangement of the DNA-binding domains that would ultimately cause the reverse phenotype. The V99E mutation disrupts the repressor’s structure, making it unable to bind to the operator DNA sites. It requires Anhydrotetracycline (ATc) to stabilize the structure and regain DNA-binding ability, leading ATc to act as a co-repressor rather than an inducer **(Scholz et al., 2004)**.

In addition, we have constructed integration-proficient variants of the vectors carrying the L5 integrase and its respective attP site. This allows integration of the vector at the specific attB site on the bacterial genome, providing more controlled regulation of the cloned genes **(Arnold et al., 2018)**.

The library’s design thus capitalizes on lessons from prior systems, including the dynamic range of TetR-regulated promoters, the stability of L5 attB integration and low gene expression levels of an integrative vector.

## Results

### Construction of pMT2 library of inducible vectors with *E. coli* TetRO system and polypeptide tags

The construction of a plasmid library incorporating the tetracycline-inducible promoter from *Corynebacterium glutamicum* was previously reported from our laboratory **(Seniya et al., 2020)**. However, these plasmids, known as the pMT series, exhibited significant leaky expression in *M. smegmatis*, limiting their application in tightly controlled gene expression and the stable maintenance of toxic genes. To address this limitation, we here present a second-generation vector library, designated pMT2, that incorporates the *E. coli tet* regulatory system and is optimized for transcriptional regulation in mycobacteria **(Meijers et al., 2020)**.

The *E. coli tet* regulatory system was first adapted for mycobacteria by incorporating the *E. coli Tn10* operator into a strong mycobacterial promoter. A 17-nucleotide operator sequence was inserted immediately downstream and two nucleotides upstream of the -35 site within the mycobacterial P_*smyc*_ promoter. The Tet repressor (*tetR*) was placed under the control of the weaker constitutive P_*imyc*_ promoter **(Ehrt et al., 2005)**. While this system demonstrated a strong induction in response to its cognate inducer, Atc, it also exhibited significant basal expression in the absence of the inducer, hindering the cloning of toxic proteins under this promoter. This issue was addressed by further optimizing the promoter by repositioning the operator from upstream of the -35 site to downstream of the transcription start site and incorporating the *rrnB*-UP element upstream of the -35 site. Additionally, *tetR* was placed under the control of the strong mycobacterial P*smyc* promoter. To further minimize leaky expression, the construct also includes a *rrnB* terminator and a bidirectional transcription terminator upstream of pTet to prevent transcriptional read-through from upstream regions. These modifications significantly reduced the promoter’s leaky expression, as demonstrated by the successful stable transformation of CRISPR-Cas9 constructs targeting essential genes **(Rock et al., 2017)**. Given the reported performance enhancement, we selected the optimized promoter sequence for use in our constructs.

The optimized Tet promoter sequence, along with the gene encoding the repressor protein and its promoter, was obtained from pCRISPR-Sth-Cas9-L5 (Addgene plasmid #140993). The Tet promoter cassette includes an NdeI site between the Tet promoter and the upstream transcription terminator sequence. To enable subsequent cloning using a different NdeI site, this site was first disrupted by NdeI digestion, followed by end-repair and ligation. Following this modification, the Tet promoter and repressor cassette were amplified separately from the NdeI-disrupted pCRISPR plasmid. The primers were designed with partial overlaps to enable fragment assembly via overlap extension PCR, resulting in a 1167 bp product. To minimize potential interference from the Psmyc promoter driving *tetR* on the Tet-regulated promoter, the fragments were assembled in diverging orientations.

The primers were synthesized with additional sequences to append a NotI and NdeI site at either end of the overlapped PCR product. These same restriction sites flanked the Tet promoter-repressor cassette in the pMT-His plasmid. The plasmid was digested with NotI and NdeI, allowing the insertion of the Tet promoter-repressor cassette at the corresponding sites, resulting in the construction of pMT2_His.

The nucleotide sequence of the inserted cassette was confirmed by Sanger sequencing. Subsequently, the NotI-NdeI cassette containing the new TetRO system was excised from pMT2_His and subcloned into the pMT-GFP, pMT-Flag, and pMT-GST vectors, replacing their existing TetRO cassettes. This resulted in the generation of pMT2-GFP, pMT2-Flag, and pMT2-GST, respectively.

As a shuttle vector, pMT2 is provided with a ColE1 *E. coli* (*oriE*) and mycobacterial origin of replication, *oriM*, which allows it to replicate in both *E. coli* and mycobacteria. The pMT2 vector provides two restriction sites (HpaI and EcoRV) for cloning of genes that will result in N-terminally or C-terminally tagged proteins. The vector also provides a Kanamycin resistance gene coding for an aminoglycoside phosphotransferase as a selectable marker.

### Construction of pMRT2 library of repressible vectors with a Reverse Tetracycline repressor (RevTetR) and polypeptide tags

We used the pMT2 vector as the template to construct the tetracycline-repressible vector, pMRT2. Several RevTetR mutants ranging from single to multiple mutations have previously been described **(Kamionka et al., 2004; Scholz et al., 2004)**, of which we selected two mutants, V99E, and E15A / L17G / L25V, based on their minimal mutation profiles, coupled with favorable expression and repression levels upon the addition of ATc.

The vector was made by reversing the Tetracycline repressor’s (TetR) activity by substituting the valine at the 99th position for glutamic acid (V99E) for the first mutant (RevTetR_E). The substitution was done through site-directed mutagenesis using the primers listed in Table 1. Meanwhile the second mutant (RevTetR_AGV) was made by mutating three amino acids thus achieving E15A, L17G, and L25V. Upon further testing, we observed that the RevTetR_AGV mutant exhibited higher GFP expression than the RevTetR_E mutant in the absence of ATc. However, the RevTetR_AGV mutant did not exhibit a responsive repression upon the addition of ATc, even at higher concentrations. Therefore, we chose to proceed with the RevTetR_E mutant, which demonstrated adequate GFP expression coupled with effective repression upon the addition of ATc. The constructed pMRT2 plasmid thus contains the RevTetR_E, henceforth referred to as RevTetR, which in the absence of ATc cannot bind to the TetO, leading to the expression of downstream gene. In contrast, when ATc is added, it binds to the RevTetR, and the complex binds to the TetO, thereby repressing the downstream gene expression. The different tags for pMRT2 were constructed similarly using the corresponding pMT2 tagged vector as the template.

**Table 1:**
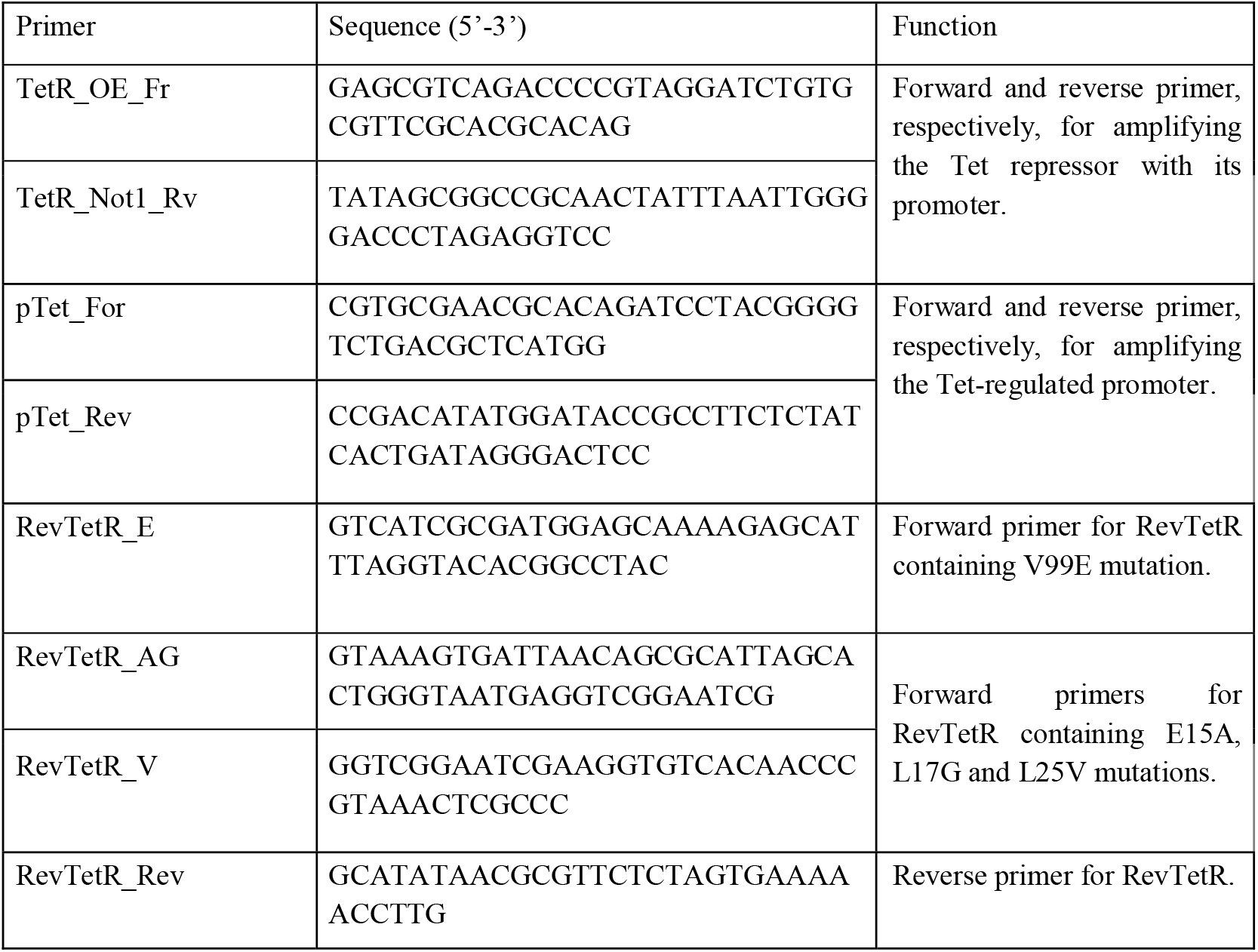
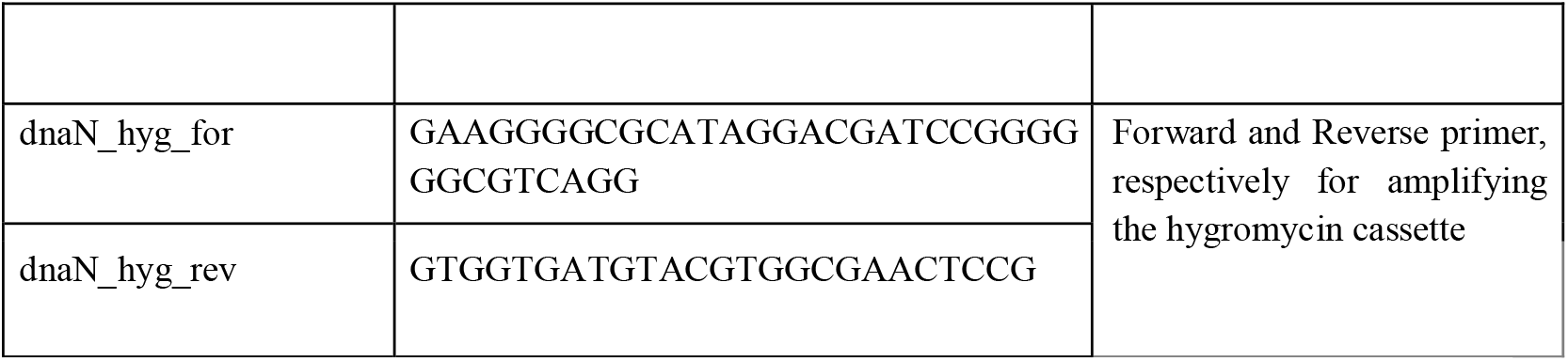
List of primers used in the present study. Sequence of the primer is given. Function corresponds to the use of the primer for a specific purpose.

### Construction of integrative Tetracycline inducible and repressible vectors and polypeptide tags

The pMT2 and pMRT2 plasmids contain the pAL5000 vector-derived mycobacterial origin of replication and exist as episomal plasmids that maintain an average copy number of 5 in *M. smegmatis* **(Ranes et al., 1990)** We further expanded our library by constructing a series of integrative vectors designed for stable insertion into the mycobacterial genome, enabling single-copy integration that remains stable in the absence of a selection marker. The integrative vector backbone, designated pMIT2, was constructed by replacing the *oriM* in the pMT2 backbone with the integrase gene cassette and *attP* integration site from mycobacteriophage L5. The integrase gene and *attP* site were isolated from the previously reported pMIA vector **; (Gangakhedkar & Jain, 2024; Seniya & Jain, 2022)** by digesting the vector with NcoI and SpeI. The *oriM* was removed from the pMT2 vectors using the same enzymes, and the integrase-*attp* cassette was inserted. Thus the vectors pMT2-His, pMT2-GFP, pMT2-Flag and pMT2-GST vectors gave rise to pMIT2-His, pMIT2-GFP, pMIT2-Flag, and pMIT2-GST, respectively in the integrative design. Subsequently, pMIRT2 versions of these vectors viz. pMIRT2-His, pMIRT2-GFP, pMIRT2-Flag, and pMIRT2-GST were generated from the respective pMIT2 vectors by converting TetR into RevTetR through site-directed mutagenesis as described above. Additionally, a modified variant of the integrative vectors replacing the kanamycin resistance cassette with the hygromycin resistant cassette was created. The pMIT2 and the pMIRT2 vector libraries coding for hygromycin resistance were labelled, pANTet and pANRTet, respectively.

### Characterization of promoter response to anhydrotetracycline in mycobacteria

The promoter activity in the newly constructed vectors was monitored through a time-based GFP fluorescence assay against the pMT-GFP vector in *M. smegmatis*. GFP fluorescence was measured to quantify the level of gene expression in the presence and absence of anhydrotetracycline (Atc). The pMT2-GFP and pMIT2-GFP showed excellent gene expression as seen from the high fluorescence upon induction by ATc with negligible leaky expression in the absence of ATc (Fig 3B-C). This when compared to poor gene expression and high leaky expression in pMT_GFP (Fig 3A) points to the exceptional regulatory capability of the *E. coli*-derived TetRO system present within the pMT2 and pMIT2 vectors. On the other hand, the repressible vectors pMRT2 and pMIRT2 showed remarkable repression of the expression upon addition of 100ng/ml ATc.; higher gene repression was observed at higher ATc concentration of 300 ng/ml, indicating that the repression by RevTetR is ATc concentration dependent (Fig 3D-G).

**Fig 1:**
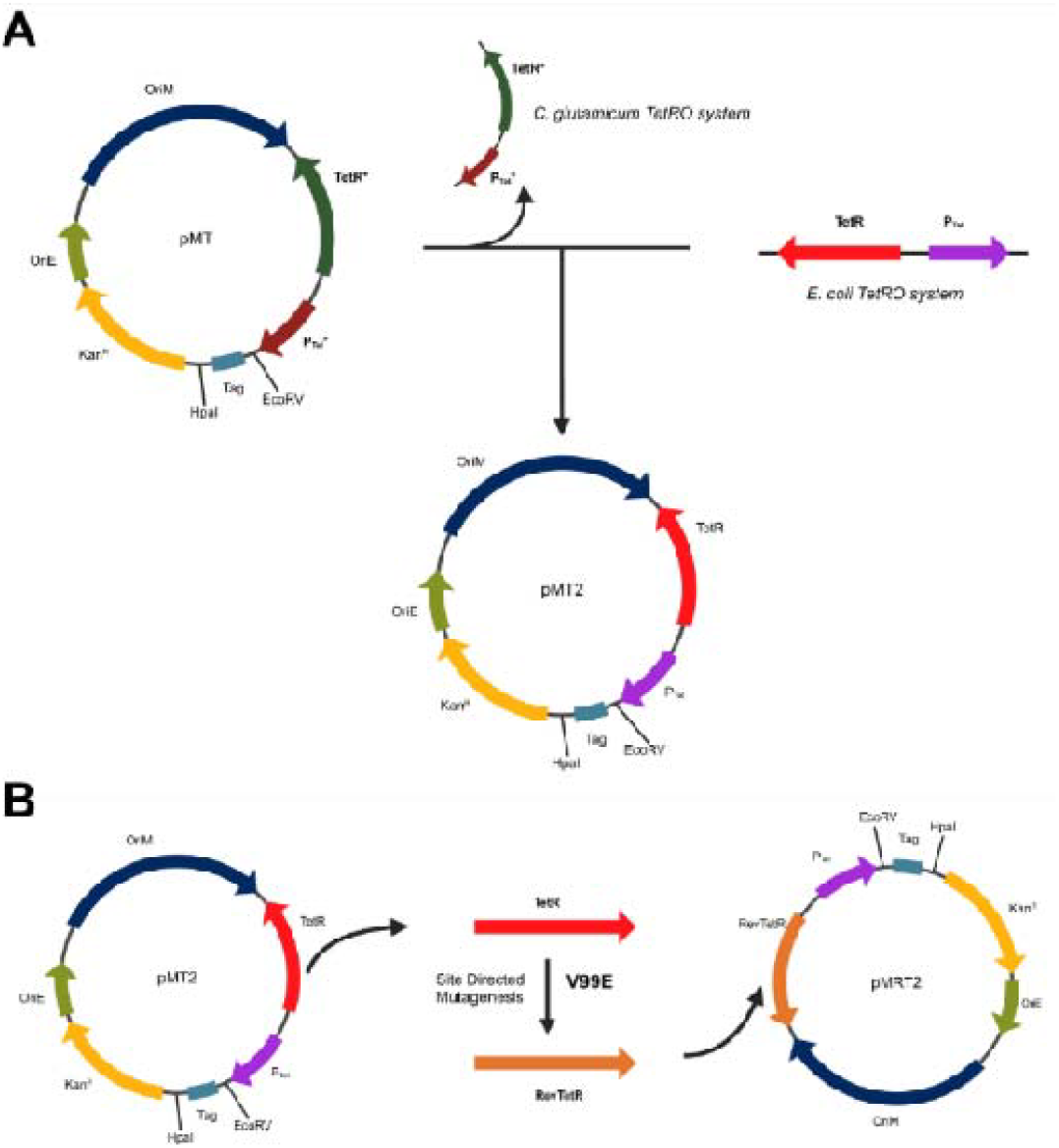
Construction of the Tetracycline inducible and repressible vector library. **(A)** Construction of pMT2 vector library using an *E. coli* Tet-regulation system. To enhance transcriptional control, the original *C. glutamicum* Tet regulatory module in the pMT vector was replaced with the *E. coli* Tet Repressor-Operator system. The resulting pMT2 library retains the backbone of pMT while yielding tighter repression in the absence of the inducer. (B) Construction of the pMRT2 vector library employing a reverse Tet repressor (RevTetR). A mutation was introduced into the Tet repressor (TetR) in pMT2 vector, generating a reverse TetR (RevTetR) that exhibits opposite regulatory behavior compared to wild-type. In this configuration, the addition of anhydrotetracycline (ATc) promotes RevTetR binding to the *tet* operator (*tetO*), resulting in repression, which, in contrast to wild-type TetR, dissociates from *tetO* upon ATc binding. The resulting Tetracycline repressible vector is labelled as pMRT2.

**Fig 2:**
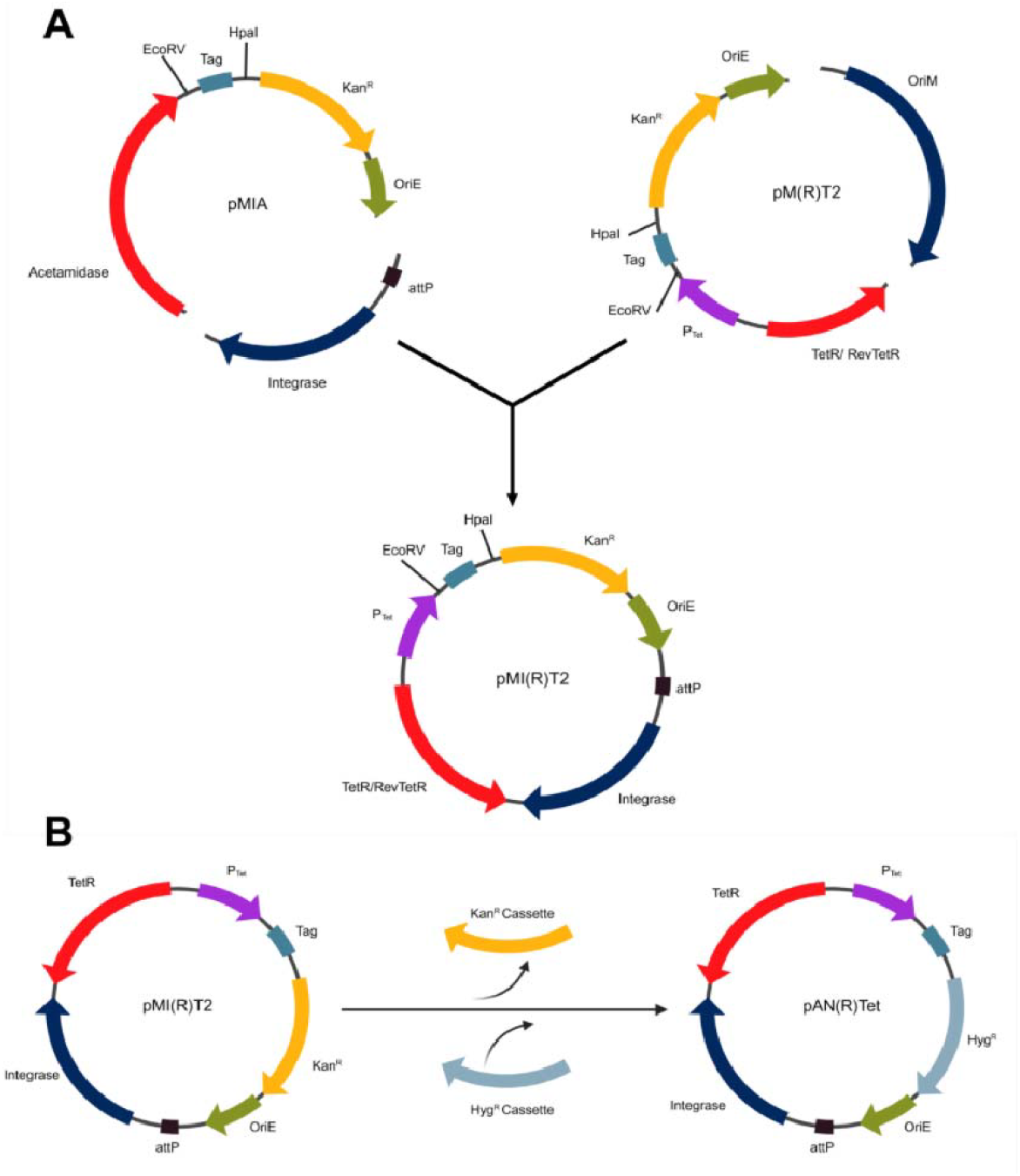
Construction of the Integrative Tetracycline inducible and repressible vector libraries. **A)** To convert the episomal pMT2 and pMRT2 vectors into single-copy integrating system, the *oriM* in the multi-copy plasmid was replaced with the Mycobacteriophage L5 integrase cassette and its cognate phage attachment site (*attP*) sourced from the pMIA vector. This yielded the pMIT2 and pMIRT2 vector libraries .**B)** A variant of pMIT2 and pMIRT2 vector libraries was constructed by replacing the kanamycin-resistance cassette with the hygromycin-resistance cassette derived from pVV16 and labelled as pANTet and pANRTet, respectively.

**Fig 3:**
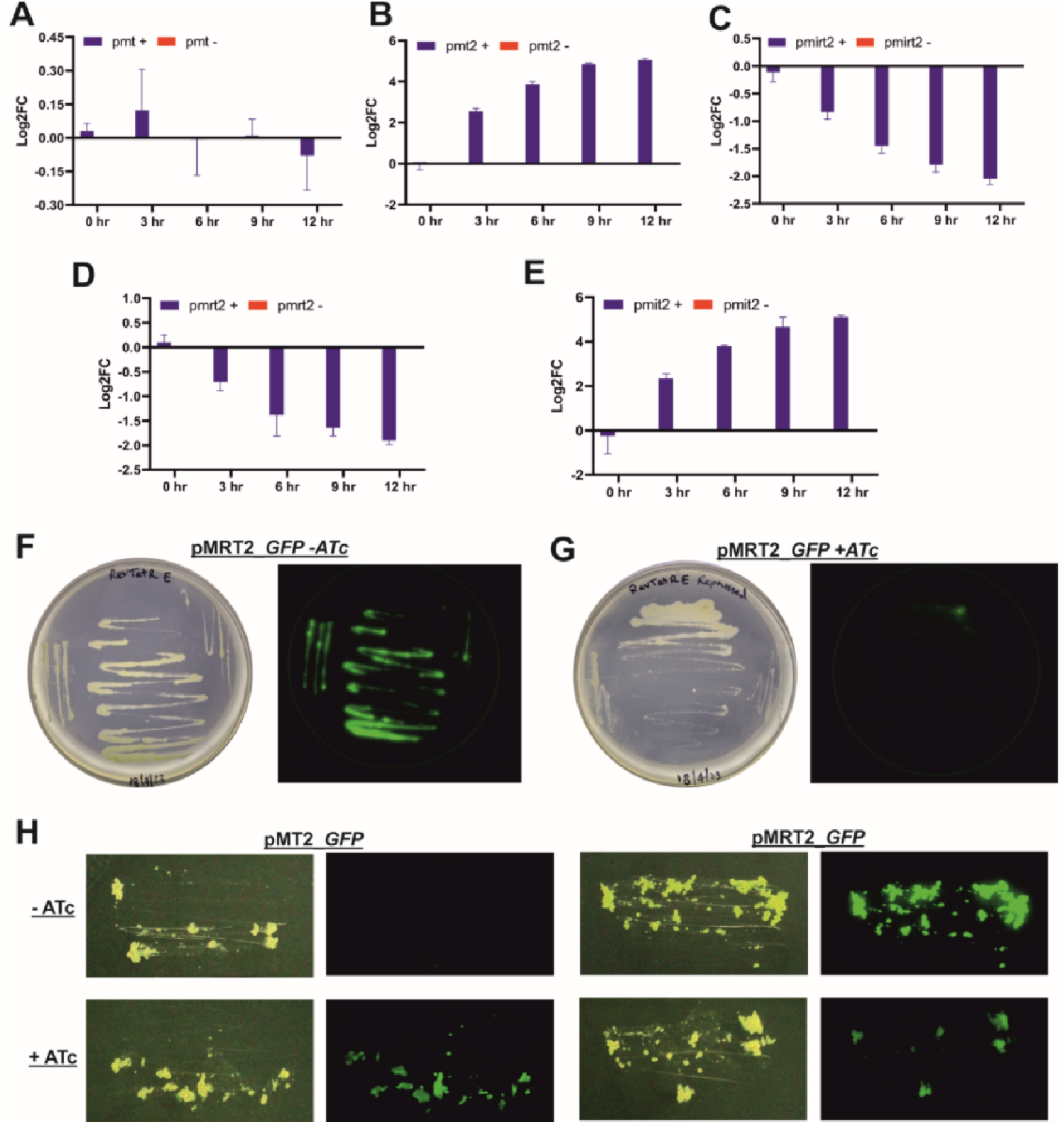
Characterization of the promoter activity. GFP fluorescence was measured over time from cells carrying different vectors in the presence and absence of ATc to determine the promoter activity. Log2FC was calculated using the -ATc sample as reference for **(A)** pMT-GFP, **(B)** pMT2-GFP, **(C)** pMIT2-GFP, **(D)** pMRT2-GFP, **(E)** pMIRT2-GFP in *M. smegmatis*. GFP repression in pMRT2-GFP carrying-M. smegmatis in the absence (F) and presence (G) of ATc. **(H)** GFP expression was tested in *M. bovis* BCG cells carrying pMT2 and pMRT2 vectors in the presence (+) and absence (-) of ATc.

**Fig 4:**
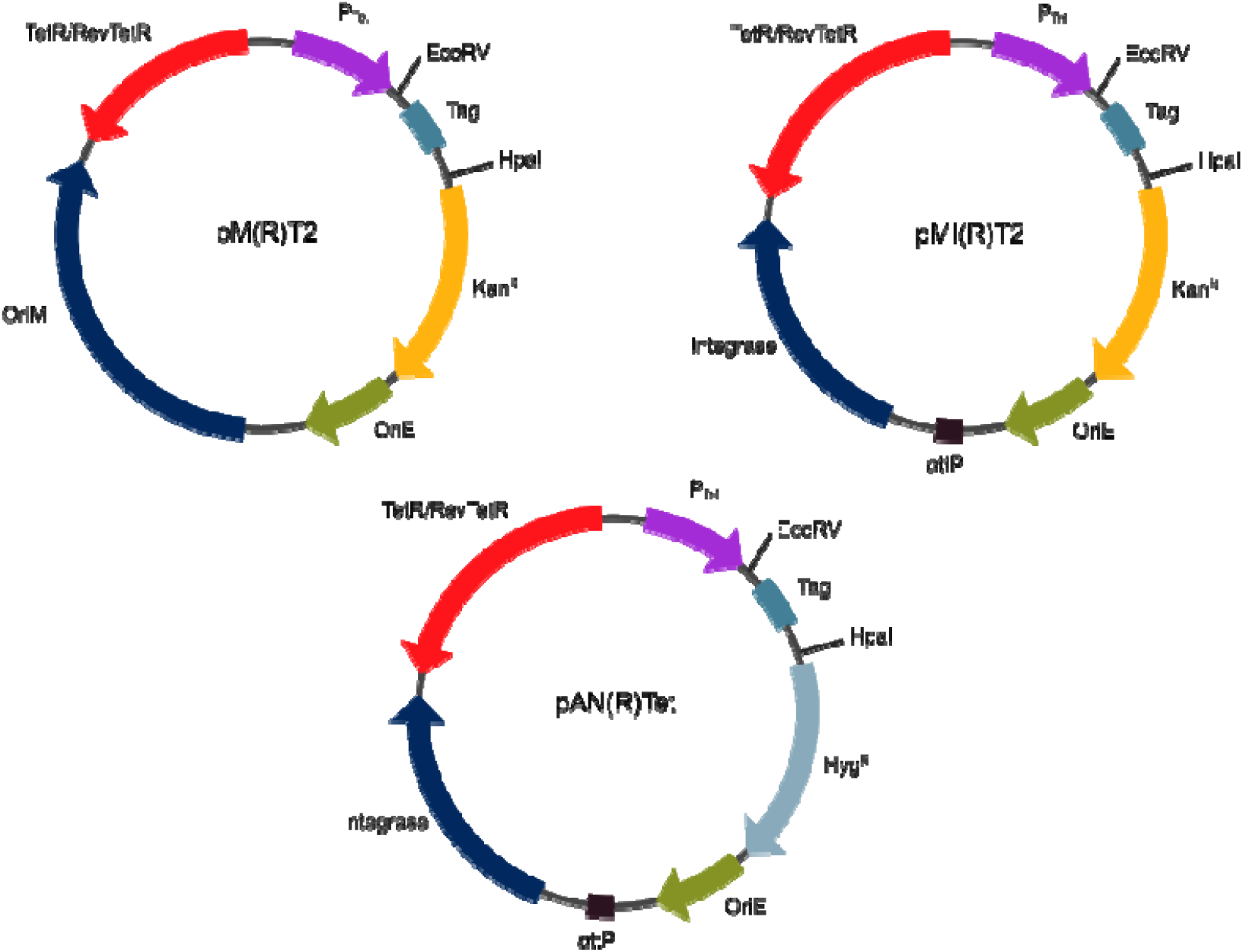
Schematic representation of the proposed vector libraries. The episomal vector libraries are labelled as pMT2 containing TetR and pMRT2 containing RevTetR. The integrative vector libraries are labelled as pMIT2 containing TetR and pMIRT2 containing RevTetR. The vectors are provided with EcoRV and HpaI restriction sites for cloning where the cloned gene can be tagged with one of four tags (viz. 6xHis, GFP, GST or Flag). The vectors carry the Kanamycin-resistance cassette (or hygromycin cassette in case of pANTet and pANRTet) along with a mycobacterial high-copy number OriM (for episomal vectors) or L5 integrase cassette and attP site (for integrative vectors).

Next, we studied the gene expression and regulation from pMT2 and pMRT2 vectors in *M. bovis* BCG. We observed excellent induction of GFP in pMT2 along with good GFP repression in pMRT2 in the presence of ATc. Although GFP expression levels in pMT2 were not as high as the unrepressed pMRT2 vector even with 300 ng/ml of ATc, we believe that for slow growing mycobacteria such as BCG, higher expression can be achieved with higher concentrations of ATc. Overall, the data shows that our vector library is compatible with BCG with good gene expression and regulation.

## Discussion

The rise of mycobacterial diseases, including tuberculosis and recently non-tuberculous mycobacterial (NTM) infections that are becoming harder to treat due to drug resistance, has burdened the medical industry and patients alike. This calls for more research into the biology of mycobacteria, which also means the need for more tools to study these bacteria. Our constructed mycobacterial vectors present here provide tool to further mycobacterial research. As a highly studied and robust regulatory system, the Tetracycline regulation system was selected for these vectors in addition to improving our laboratory’s previously constructed pMT vector library. The switch from the Corynebacterial Tet system used in pMT to the more regulated E. coli Tet system and modifications to the Tet-regulated promoter in pMT2 led to higher expression levels during induction with minimal leaky expression. The E. coli Tet system was taken from the pCRISPRx-Sth1-Cas9-L5 vector. Along with the construction of an improved pMT2 vector library, we have developed a repressible counterpart, the pMRT2 vector library. The pMRT2 vector was constructed by reversing the phenotype of TetR, which was achieved by introducing specific mutations within TetR, particularly the V99E mutation.

We also present here the integrative variants for pMT2 and pMRT2 vectors. The integrative vector use the mycobacteriophage L5 integrase and attP site obtained from the previously described pMIA vector and can integrate into the genome of mycobacteria at the respective attB site. Integrative vectors will allow for the study of gene interactions through co-expression studies, providing more insights into bacterial biology. The pAN(R)Tet series substitutes the Kanamycin resistance marker with the hygromycin in the pMI(R)T2 series. All the vectors constructed in this study are also designed to carry different tags for the cloned genes, including 6x His, FLAG, GFP, and GST tags.

The pMT2 and pMRT2 vectors also show good expression and regulation in M. bovis BCG strain, which expands their use in clinically important mycobacterial strains, especially towards vaccine development.

## Materials and methods

### Plasmids, bacterial strains, media, and growth conditions

pCRISPRx-Sth1-Cas9-L5 vector was obtained from Hatfull, while the pMT expression vector was previously developed in the lab. Cloning and screening were done in E. coli strain XL1-Blue (Stratagene), grown in LB broth (Difco) at 37°C supplemented with 50 μg/ml kanamycin (MP Bio) and constant shaking at 200 rpm or on LB agar plates supplemented with 50 μg/ml kanamycin. *M. bovis* BCG strain (BCG) obtained from Ramandeep Singh and *M. smegmatis* mc^2^155 (Msm) was used for the mycobacterial expression studies and grown in MB 7H9 (Difco) media supplemented either with 2% (w/v) glucose and 0.05% (v/v) Tween80 for Msm or 0.2% (v/v) glycerol and 0.05% Tween80 for BCG. MB 7H9 agar plates were supplemented with either 2% (w/v) glucose for Msm or 0.5% (v/v) glycerol and 4% (v/v) OADC (Oleic acid, Albumin, Dextrose, Catalase) (Difco) supplement for BCG.

Plasmids generated in this study (Table 1) were electroporated into *M. smegmati*s, and the bacteria were selected on the required antibiotic. Bacteria were incubated at 37°C with constant shaking at 200 rpm. Where necessary, 50 μg/ml of kanamycin or hygromycin (MP Bio) and 100ng/ml ATc (Sigma) was used for mycobacterial culture.

### Molecular Cloning

Restriction enzymes, Antarctic phosphatase, T4 polynucleotide kinase, T4 DNA ligase, etc., were procured from New England Biolabs. All the enzymes were used as per the manufacturer’s instructions. Fast DNA End Repair Kit was purchased from Thermo Fisher Scientific. DNA oligos were obtained from Thermo Fisher Scientific. The Plasmid prep kit and Gel extraction kit were purchased from Favorgen and used according to the manufacturer’s instructions.

### PCR and Site-directed mutagenesis (SDM)

Q5 polymerase, DpnI were procured for New England Biolabs and used as per the manufacturer’s instructions. Q5 polymerase was used in all PCR and SDM reactions using oligos listed in Table 1, while DreamTaq (Thermo) was used for colony PCR (cPCR). SDM reaction was performed as described elsewhere **(Singh et al., 2013; Dubey et al., 2016)**

### Promoter activity

Cultures of Msm cells harbouring the pMT, pMT2 and pMRT2 GFP plasmids were grown till OD = 0.6 and 100ng/ul ATc was added to the samples and were collected at different timepoints. The collected samples were added to 96 well plate and the absorbance at 600nm and fluorescence at Ex = 495nm and Em= 525nm were recorded using spectramax M5 plate reader (molecular devices). The fluorescence per unit absorbance was calculated and the log2fold change between the +ATc and –ATc samples were plotted.

The assay was repeated for BCG cells harbouring pMT2 and pMRT2 GFP plasmids. Cultures were grown till OD = 0.2 and 300ng/ml ATc was added. Samples were collected every 24 hours and absorbance and fluorescence was measured and log2fold change was calculated and plotted Data was plotted using Graphpad Prims v8.0.

### Fluorometric plate scanning

Colonies of Msm and BCG strains were patched onto MB 7H9 agar plates supplemented with glucose or glycerol and kanamycin. ATc at a concentration of 100 ng/mL was added to the plates as needed. The plates were incubated at 37°C and the patches were allowed to grow. The plates were scanned using a Typhoon FLA 9000 Biomolecular Imager (GE) at 488nm with the EGFP filter. Successive images of the plates were taken with an EOS 1500D (Canon) camera, showing the growth of the patches.

## Notes

### Competing Interest Statement

The authors have declared no competing interest.

